# Comparative transcriptomics of Venus flytrap (*Dionaea muscipula*) across stages of prey capture and digestion

**DOI:** 10.1101/2023.09.21.558761

**Authors:** Jeremy D. Rentsch, Summer Rose Blanco, James H. Leebens-Mack

**Affiliations:** Department of Biology, Francis Marion University, Florence, SC, USA; Department of Plant Biology, University of Georgia, Athens, GA, USA

**Keywords:** Dionaea muscipula, Venus flytrap, Transcriptomics, Botanical Carnivory

## Abstract

The Venus flytrap, *Dionaea muscipula*, is perhaps the world’s best-known botanical carnivore. The act of prey capture and digestion along with its rapidly closing, charismatic traps make this species a compelling model for studying the evolution and fundamental biology of carnivorous plants. There is a growing body of research on the genome, transcriptome, and digestome of *Dionaea muscipula*, but surprisingly limited information on changes in trap transcript abundance over time since feeding. Here we present the results of a comparative transcriptomics project exploring the transcriptomic changes across seven timepoints in a 72-hour time series of prey digestion and three timepoints directly comparing triggered traps with and without prey items. We document a dynamic response to prey capture including changes in abundance of transcripts with Gene Ontology (GO) annotations related to digestion and nutrient uptake. Comparisons of traps with and without prey documented 174 significantly differentially expressed genes at 1 hour after triggering and 151 genes with significantly different abundances at 24 hours. Approximately 50% of annotated protein-coding genes in Venus flytrap genome exhibit change (10041 of 21135) in transcript abundance following prey capture. Whereas peak abundance for most of these genes was observed within 3 hours, an expression cluster of 3009 genes exhibited continuously increasing abundance over the 72-hour sampling period, and transcript for these genes with GO annotation terms including both catabolism and nutrient transport may continue to accumulate beyond 72 hours.

## 1 Introduction

Charles Darwin described the Venus flytrap, *Dionaea muscipula* Ellis, as both “one of the most wonderful [plants] in the world” and a “horrid prison with closing walls” (Darwin, 1875). The Venus flytrap is perhaps best known for its active, snapping trap mechanism that it uses to capture prey for nutrition. This snap trap mechanism is shared with the related aquatic waterwheel plant, *Aldrovanda vesiculosa* L. (Droceraceae) (Cameron et al., 2002), implying the origin of the snap trap in a common ancestor. Droseraceae and three other plant families with carnivorous species - Drosophyllaceae, Nepenthaceae, Dioncophyllaceae - form a clade within the Caryophyllales along with the non-carnivorous family Ancistrocladaceae indicating an ancient origin of carnivory, and subsequent losses of carnivory in Ancistrocladaceae and two of three genera within Dioncophyllaceae (Heubl et al., 2006; Renner and Specht, 2011). Carnivory has evolved independently more than five times in angiosperm history (Givnish 2014, Lin et al. 2021) as an adaptation to thrive in nutrient-deficient soils in sunny, wet habitats (Givnish et al., 1984). The Venus flytrap is no exception to this rule, growing in semi-pocosins or semi-savannah areas (Roberts and Oosting, 1958).

The Venus flytrap closes in response to the stimulation of mechanosensing trigger hairs, which can be found on the adaxial side of the trap (Darwin 1875). Stimulation of these hairs in rapid succession causes a very rapid shift in action potentials, two of which triggers trap closure (Burdon-Sanderson, 1873; Jacobson, 1965; Volkov et al., 2007, Hedrich and Kreuzer, 2023). Once a trap accumulates sufficient electrical charge from these action potentials, proton transport (Williams and Bennett, 1982; Rea, 2006), and ATP hydrolysis begin (Jaffe, 1973), aquaporin channels open (Volkov et al., 2008), and water transport causes a rapid change in turgor across the trap (Hodick and Sievers, 1986). Successful trap closure begins with increased trap turgor followed by a simultaneous expansion of the outer epidermis and a shrinkage of the inner epidermis (Brown, 1916; Sachse et al., 2020), changing the overall trap curvature from convex to concave, resulting in trap closure (Markin et al., 2008). Upon properly sealing, a cocktail of digestive fluids is released into the interior of the trap and prey digestion takes place.

Over recent years, an array of digestive enzymes in the Venus flytrap have been well characterized. These digestive enzymes include a cysteine endopeptidase named dionain (Takahashi et al., 2011), as well as peroxidases, nucleases, phosphatases, phospholipases, a glucanase, chitinases, aspartic proteases, and a serine carboxypeptidase (Schulze et al., 2012). The initial release of digestive fluids is triggered by the action potentials generated from mechanical stimulation (Ueda et al., 2010; Volkov et al., 2013b). These action potentials are stimulated until prey death and are important in the early establishment of the digestive cycle (Volkov et al., 2013a).

After prey death, the trap may require chemical feedback from partially digested prey to complete the release of digestive enzymes and promote nutrient absorption and transport. In aseptically grown *Drosera rotundifolia* L., for example, the application of crustacean chitin to the carnivorous plant leaves induced a marked increase in chitinase activity (Matusíková et al., 2005), demonstrating the link between enzyme release with the addition of chemical cues. Jakšová and others (2020) showed that digestive enzyme regulation is not substrate specific, with bovine serum albumin (BSA) protein eliciting an upregulation of proteolytic enzymes, phosphatases, and chitinases.

At the same time, the trap absorbs nutrients – typically the domain of roots. After capture, the prey is digested, releasing amino acids and peptides. Glutamine is deaminated to produce ammonium ion (NH_4_^+^) which is then channeled into gland cells via the upregulated ammonium channel DmAMT1 (Scherzer et al., 2013). Venus flytraps also use prey-derived amino acids as a substrate for cellular respiration (Fasbender et al., 2017) even with an abundance of atmospheric CO_2_. In the carnivorous plant genus *Nepenthes*, transporters for amino acids and peptides are also expressed in the modified leaves (Schulze et al., 1999).

Hormonal signaling also contributes to prey capture and digestion in carnivorous plants. The phytohormone jasmonic acid (JA) plays a critical role in both localized and systemic plant responses to injury (as reviewed in Wasternack and Hause, 2013) through the synthesis of secondary metabolites (Pauwels et al., 2009) or by invoking changes in gene expression and growth form (Yoshida et al., 2009). In *Lycoris aurea* (L’Hér.) Herb., for example, treatment of seedlings with methyl jasmonate resulted in 4,175 differentially expressed genes when compared to control plants (De Geyter et al., 2012). Venus flytraps and other species in the carnivorous clade of the Caryophyllales also utilize the JA pathway to elicit prey capture and digestion responses (Nakamura et al., 2013; Yilamujiang et al., 2016; Pavlovič et al., 2017).

Recently, a significant amount of work has gone into understanding the genomes of convergently evolved carnivorous plant lineages, including that of the Venus flytrap. The genome of *Utricularia gibba* L., a carnivorous bladderwort, was found to have a haploid size of just 82 MBP/C in size while still encoding 28,500 genes (Ibarra-Laclette et al., 2011), comparable to the number of genes in *Arabidopsis thaliana* (Pasha et al., 2020). *Genlisea aurea* A.St.-Hill in the Lentibulariaceae is smaller still, with a nuclear genome size of only 64.6 MBP/C. These nuclear genomes are the two smallest vascular plant genomes sequenced to date.

Palfalvi and others (2020) sequenced the genomes of three carnivorous plants in the Droseraceae, *Dionaea muscipula, Drosera spatulata* Labill., and *Aldrovanda vesiculosa* L. with genome sizes of 3,187 MBP/C, 293 MBP/C, and 509 MBP/C, respectively. The authors point to a genome duplication event in an ancestral Droseraceae species and an additional genome duplication event in the ancestor of *A. vesiculosa*. Venus flytrap has a much larger genome size than *Drosera* or *Aldrovanda* due to a massive expansion of long terminal repeat (LTR) retrotransposons, which comprise at least 38.78% of its genome. Palfalvi et al. (2020) inferred a reduction in the number of protein-coding genes in the ancestral Droseraceae lineage, but interestingly, gene families with functions associated with prey attraction, nutrient uptake, and digestion increased in size. Here, we investigate change in transcript abundance over a time course following prey capture in Venus flytrap. We compare transcripts from traps that were triggered with and without prey and characterize sets of genes exhibiting similar temporal changes in transcript abundance over a 72-hour time course following prey capture. Our findings document a complex and dynamic transcriptional process with significant differences between triggered traps with and without prey seen in transcript profiles just one hour after closure.

## 2 Materials and Methods

### Plant care and total RNA isolation

Seed grown plants obtained from tissue culture stock provided by Michael Kane (University of Florida) were grown in an environmental chamber maintained at 22°C supplemented with fluorescent lights which were cycled 16 hours on and 8 hours off a day. Plants were grown in a 1:1 mix of peat moss and perlite and watered with distilled water at under 50 parts per million of total dissolved solids. Experimental traps were fed with live darkling beetle (Tenebrionidae) larvae. Prior to RNA isolation, traps were harvested with sterile scissors just distal to the point of petiole attachment. Tissue was harvested from plants grown from seed that was derived from a single plant grown from tissue culture (tissue culture provided by Michael Kane, University of Florida). A single trap was harvested from each of the 44 plants randomly assigned to ‘prey’ and ‘no prey’ treatments before triggering trap closure. Traps without prey were sampled at 0m, 5m, 60m, and 1440m following mechanical triggering, and traps with prey were sampled at 5m, 30m, 60m, 720m, 1440m, 2880m, and 4320m after triggering trap closure. Immediately prior to harvesting, traps with beetle larvae were cut longitudinally along one side and the prey item was extracted with sterile forceps. Traps were immediately placed into liquid nitrogen for the first stage of tissue homogenization.

Bulk RNAs were isolated from each sampled trap with four biological replicates per treatment. Additionally, we isolated total RNA from petiole tissue and traps at 0 min. Total RNA was isolated using the Direct-zol kit (Zymo Research, Irvine, CA, USA) with Plant RNA reagent (Life Technologies, Carlsbad, CA, USA). One 2880m prey sample library was dropped so we only had three replicates for these two treatments.

### RNA-Seq and read processing

Illumina RNASeq library construction and RNA sequencing on an Illumina NovaSeq platform was conducted at HudsonAlpha Institute for Biotechnology. The median and minimum for read pairs after trimming are in supplemental table. Trimmomatic v0.39 (Bolger et al., 2014) was used for quality trimming and adapter clipping. Read quality was assessed using FASTQC v0.11.9 (Andrews, 2010) & MultiQC v1.8 (Ewels et al., 2016) before and after trimming. Pre- and post-trimming read counts are reported in Supplemental Table 1AC).

### Paired Timepoint Differential Expression Analysis

Trimmed reads were pseudo-aligned to the *D. muscipula* reference transcriptome index, then quantified using Kallisto v0.46.1 (Bray et al., 2016). The transcriptome index was generated using published gene models and their annotations (Palfalvi et al., 2020) downloaded from https://www.biozentrum.uni-wuerzburg.de/carnivorom/resources (*Dm_transcripts)*.

A principal component analysis (PCA) was then performed on the pseudo-aligned reads using the R software package Sleuth (Pimentel et al., 2017) to visualize the variance between sample groups and replicates. Kallisto transcript abundances were then used for pair-wise differential gene expression analyses between triggered traps with and without prey sampled at 5 min, 1 hr, & 24 hr after triggering.

Differentially expressed transcripts were then translated using SeqKit (Shen et al., 2016) and protein sequences were BLASTed against annotated *Arabidopsis thaliana* proteins (Araport11_pep_20220914) from The Arabidopsis Information Resource (Berardini et al., 2015) and the Swiss-Prot database (Bairoch & Boeckmann, 1994). GO Term Enrichment was performed using the org.At.tair.db (Carlson, 2019) and TopGO (Alexa & Rahnenfuhrer, 2023) packages in R.

### Gene Co-expression Analysis

To characterize changes in transcript abundance across sampling time points, a gene coexpression analysis was performed including Transcripts per Million (TPM) data for both treatments and all time points. The analysis was performed in R using the Simple Tidy GeneCoEx workflow (Li et al., 2023 - https://github.com/cxli233/SimpleTidy_GeneCoEx). This method detects groups of genes that display similar expression patterns. Briefly, the workflow generates a gene correlation matrix from gene expression data using the Tidyverse package (Wickham et al., 2019). Then, the matrix is used to identify genes showing variation in expression across timepoints, construct a co-expression network and detect sets (modules) with genes exhibiting highly correlated transcript abundance profiles.

Annotations of genes within each module were inspected for terms related to catabolic processes including synthesis of digestive enzymes. GO Term Enrichment for each module was performed using the org.At.tair.db (Carlson, 2019) and TopGO (Alexa, 2023) packages in R.

## 3 Results

### 3.1 Paired Time Point Differential Expression Analyses

After filtering, transcripts for 15146 and 14861 gene models were found 1 hour and 24 hours after triggering, respectively. We identified 174 and 151 differentially expressed genes between prey-fed and mechanically triggered (“no-prey”) traps at 1 hour and 24 hour time points (q-val<0.05), respectively. Interestingly, no differentially expressed genes were identified between prey-fed and no-prey traps at the 5-minute time point.

At the 1-hour time point, 103 transcripts showed significantly higher expression in prey-fed traps, while 71 transcripts showed significantly higher expression in no-prey traps. Several GO terms were enriched among DE genes at this time point including: response to stimulus, regulation of jasmonic acid mediated signaling pathway, defense response to bacterium, response to salicylic acid, and photosynthesis (Supplemental Figure 1).

At the 24-hour time point, 149 transcripts showed significantly higher expression in prey-fed traps, while just 2 transcripts showed significantly higher abundance in no-prey traps. These include a gene homologous to the flowering-time gene CONSTANS and a basic leucine zipper transcription factor. Several GO terms were enriched at this time point including response to wounding, response to jasmonic acid, defense response to bacterium, response to salicylic acid, and transmembrane transport (Supplemental Figure 1)

### 3.2 Gene Co-expression Analysis

A Principal Component Analysis (PCA) was used to visualize variation among samples and treatments. The first principal component (explaining 31.8% of the variance within the data) appears to correlate with the time (Figure 1a). As described above, comparisons of traps with and without prey did implicate differentially expressed genes at the 1- and 24-hour post-triggering timepoints, but full transcript profiles for these two treatments are overlapping in the PCA plot (Figure 1b).

**Figure 1.**
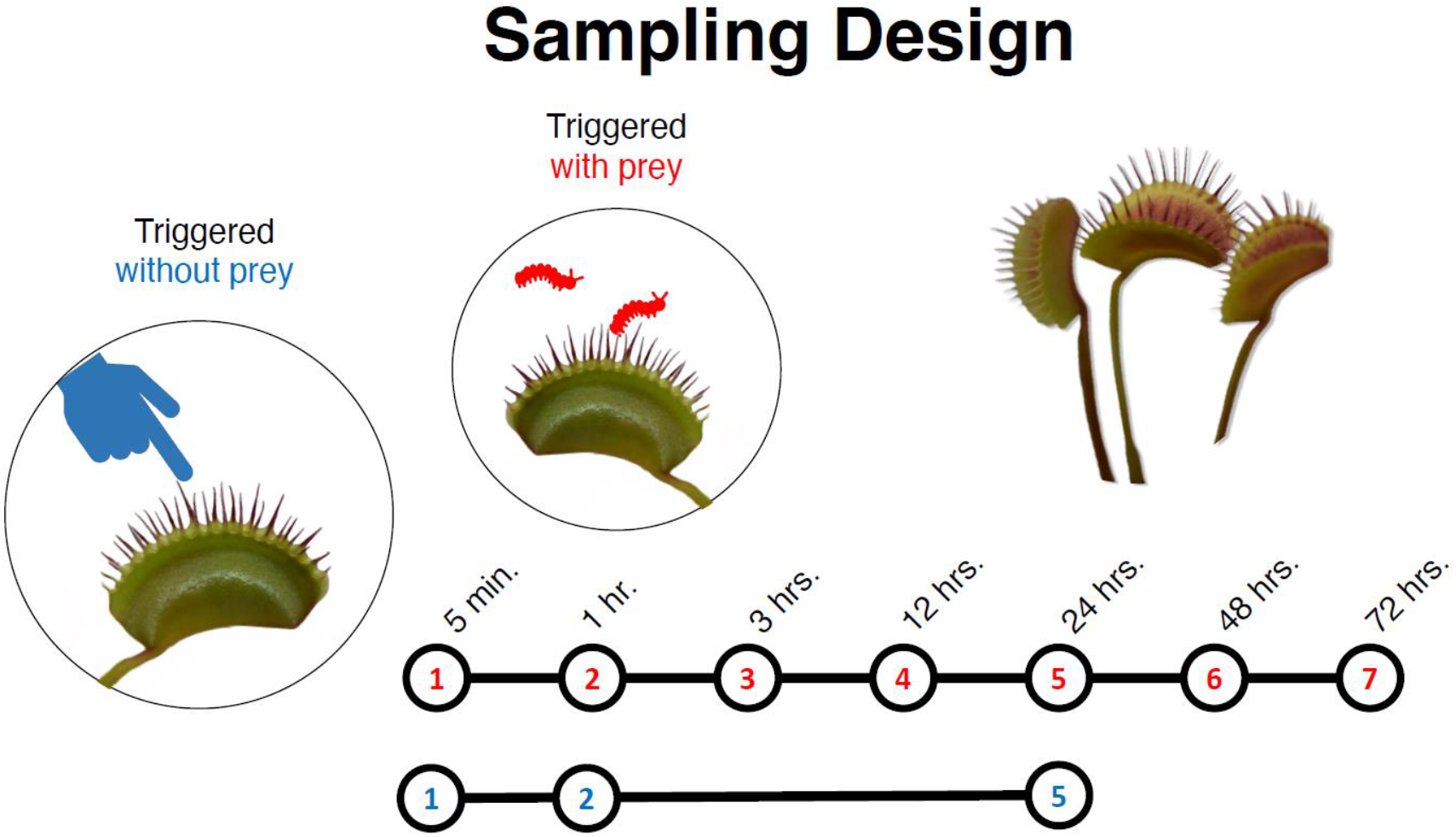
A diagram of the sampling design. Venus fly traps were either mechanically triggered (blue) or fed prey (red). Traps triggered with prey were harvested at 7 different time points between 5 minutes and 72 hours. Mechanically triggered traps were harvested at 3 different time points (5 minutes, 1 hour, and 24 hours)

Of the 14217 gene models for which abundance levels greater than five transcripts per million were observed after filtering, 10041 exhibited variation across time points. Transcript abundance profiles for these genes clustered into 14 modules, each with more than five highly coexpressed genes (Figure 3a, Supplementary Figure 2). Modules were assigned numbered names by the GeneCoEx workflow, but as is evident in Figure 3, these names are not ordered with respect to module gene number or time signature.

**Figure 2.**
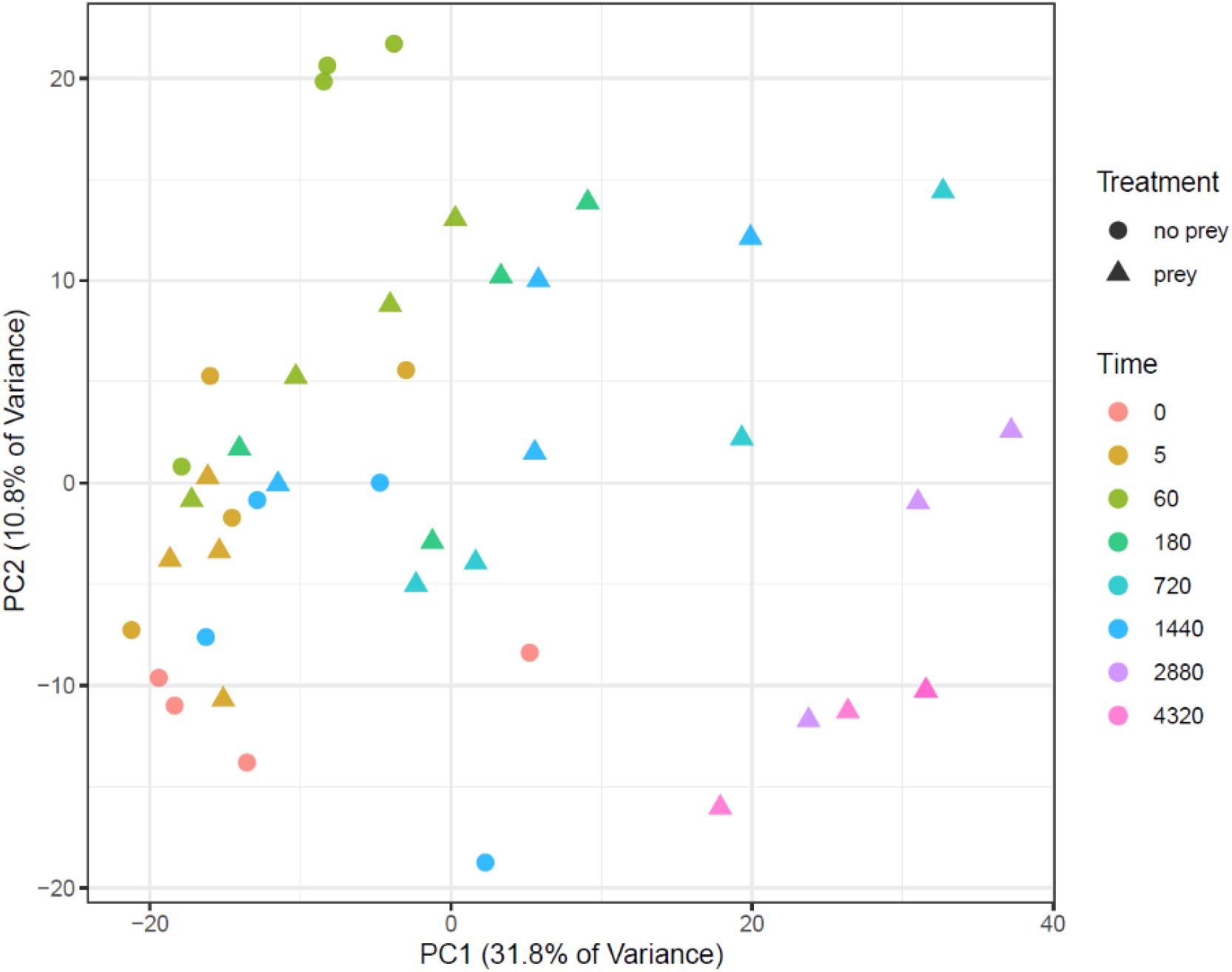
Principal component analysis (PCA) of RNA-seq data from traps. Replicates from the same time point are indicated with the same color while treatments are indicated by geometric shape (circle for mechanically triggered vs. triangle for prey-fed traps).

**Figure 3.**
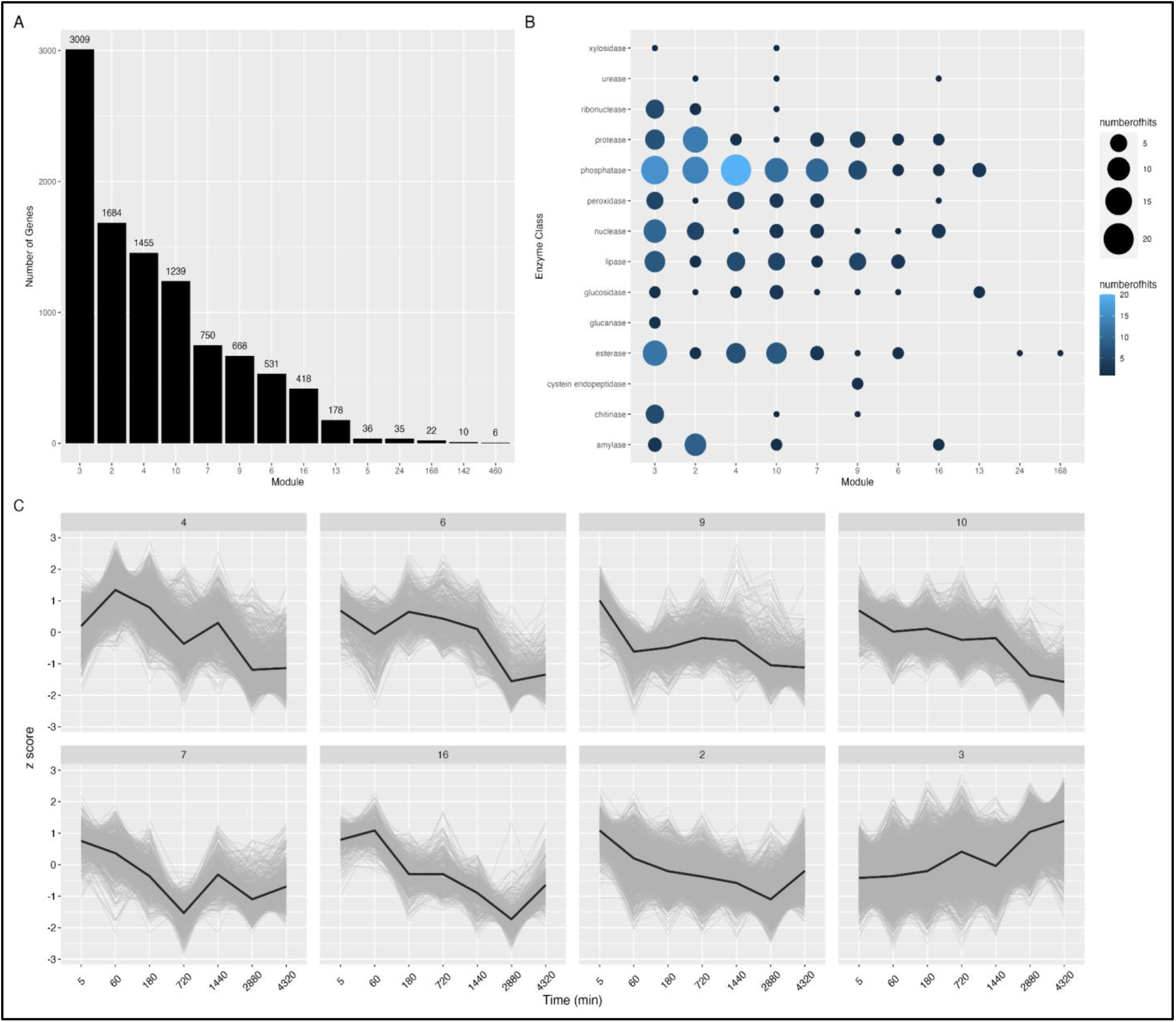
Gene Co-expression Analysis. A. Number of genes in each module with >5 highly co-expressed genes (n=14). B. The number of BLAST hit descriptions for enzymes related to digestion are shown for each module. Color and size of each point depict the number of enzyme transcripts in each module. C. Expression Z-scores for each gene in modules with >400 genes (n=8). Clusters represent co-expressed genes within each module. Mean expression is shown in the black line.

Focusing on the eight modules with more than 400 genes, Figure 3c shows that most module profiles exhibit peak transcript between 5 and 60 minutes after prey capture. Interestingly, the largest module (n=3009, Figure 3a,b module “3”; Figure 3c bottom right panel), including ∼21% of transcripts with time-structured expression, is the exception with genes showing increasing transcript abundance from 5 minutes until the last time point at 72 hours after prey capture.

Several classes of enzymes were represented in each module (Figure 3b). Phosphatases were most abundant across modules. Cysteine endopeptidases were only found in module 9. Glucanases were only found in module 3 which exhibited peak expression at 72hrs.

Additionally, many genes from the prey/no-prey DE gene lists were found among the time-signature modules. Module 3 contained 29 and 83 genes from the 1hr and 24hrs, prey/no-prey differential expression analysis, respectively. Module 4 contained 57 and 64 genes from the 1hr and 24 pair-wise differential expression analyses, respectively. Module 10 contained 35 genes from the 1hr pair-wise differential expression analysis. (Supplementary Figure 3)

## 4 Discussion

### 4.1 Transcriptomic differences between traps with prey items and ‘no-prey’ traps

It is thought that the relative rarity of botanical carnivory is, at least in part, due to the narrow range of conditions under which the habit is advantageous from a cost/benefit perspective (Givnish et al., 1984). Thus, it would be reasonable to assume that the overall fitness of the flytrap is at least partially dependent on the ability of the plant to detect failed prey capture attempts and ultimately reset the trap. It is well known that the Venus flytrap plants respond differently to triggering of traps with and without prey. The generation of the action potentials driving triggered trap closure results in a temporary decrease in photosynthetic activity to nearly zero. In traps retaining prey, the decrease in photosynthetic activity is prolonged for as long as the trap is receiving mechanical stimulation (Pavlovič et al., 2010). Therefore, triggered traps without prey temporarily lose the ability to capture new prey and experience a decrease in their photosynthetic rate without increased nutrient absorption from prey being digested.

The resetting of a falsely triggered trap is a relatively long process – on the order of several days (Volkov et al., 2011) and there are outstanding questions about what is happening at the transcriptomic level during much of this time. To that end, we examined the full transcriptomes of traps triggered with and without prey items at 5m, 60m, and 1440m (24 hrs.). Interestingly, at the five-minute time point we detect no genes being differentially expressed between traps with and without prey, suggesting the trap has not yet determined whether or not it has captured a suitable prey item. At the 60-minute time point, we identified 174 significantly differentially expressed genes between prey-fed and no-prey traps. Of the 174 DEGs, 103 genes showed higher expression in traps containing prey and 71 genes showed higher expression in traps without prey. By 60 minutes after trap closure, the annotations of transcripts exhibiting significantly higher abundance with vs. without prey imply 1) the activation of the jasmonic acid pathway and other responses to wounding, 2) a general response to bacterial and fungal pathogens, and 3) nutrient absorption.

Pathogenesis-related (PR) proteins have been described in the pitcher fluid of the carnivorous *Nepenthes alata*. The putative function of these proteins is to suppress the proliferation of putrefying bacteria on captured prey items or in the pitcher fluid (Hatano and Hamada, 2008). Unlike digestion in *Dionaea* traps, *Nepenthes* pitchers do not enclose their prey after capture, and pitchers are known to host a diverse microbiome community in their digestive fluid (Chan et al., 2016). For this reason, it may be somewhat surprising to see PR-related genes being upregulated in *Dionaea*; however, despite the sealed environment of the *Dionaea* trap, prey items may well harbor putrefying bacteria or fungi. PR proteins have been detected in the secretome of *Dionaea* previously (Schulze et al., 2012), however, the detection of PR gene transcripts after only an hour may suggest a dual role in both pathogen defense and prey digestion (Mithöfer, 2011).

Of additional interest in the realm of defense response is the abundance of transcripts for a *Dionaea* homolog of AtNPR1 (AT1G64280). AtNPR1 is a key regulator of the salicylic acid-mediated system acquired resistance pathway and is important for cross-talk with the JA pathway (Spoel et al., 2003). This regulator primes an enhanced resistance to bacterial and fungal pathogens in *Arabidopsis thaliana* (Cao et al., 1994) and *Oryza sativa* (Quilis et al., 2008). Perhaps there is a function here in the moderation between the defense response and the wounding response.

With respect to nutrient transport, a homolog to AKINBETA1 (AT5G21170) encoding a subunit of SnRK1, which acts as a positive regulation of lipid and nitrogen metabolism (Wang et al., 2020), exhibits upregulation in traps with prey just one hour after prey-capture. The expression of this subunit has been shown to increase significantly in *Arabidopsis thaliana* after treatment with ammonia nitrate(Li et al., 2009). The upregulation of AtLHT1 (AT5G40780), responsible for the uptake and cycling of amino acids and their derivatives (Hirner et al., 2006) also implies that the trap is primed for nutrient absorption only one hour after prey capture, possibly before the secretion of digestive fluid into the lumen of the closed trap.

In contrast to traps with prey, those without prey exhibit an increased abundance of transcripts for proteins to be transported to chloroplasts, perhaps contributing to the repair of chlorophyll or restoration of photosynthesis prior to the trap resetting. Among these are genes, one is involved in the production of a major leaf ferredoxin (AT1G60950), a second, ATCNFU3 (AT4G25910), is required as a molecular scaffold of ferredoxin and photosystem I (Yabe et al., 2004), and RIQ1 (AT5G08050), which encodes a thylakoid membrane protein (Yokoyama et al., 2016). Reactive oxygen species are known to spike during prey capture and digestion (Bemm et al., 2016) and evidence from other sources of oxidative stress suggests that these molecules can be particularly damaging to chlorophyll (Aro et al., 1993; Yamamoto et al., 2008), so it would not be surprising if the stress of trap closure damaged chlorophyll via the generation of reactive oxygen species.

We identified 151 significantly differentially expressed genes between prey-fed and no-prey traps at 1440 m (24 hrs.). Of the 151 DEGs, 149 genes showed higher expression in traps containing prey, while only 2 genes showed higher expression in traps without prey. Again, the 1440-minute time point is rich in genes related to plant defense and the jasmonic acid pathway. Additionally, genes related to nutrient transport continue to exhibit higher abundance in traps with prey. HAK5 (AT4G13420), for example, codes a high-affinity K+ channel that allows for the absorption of K+ from captured prey even at very low concentrations (Scherzer et al., 2015). PLT5 (AT3G18830) codes a transporter for a wide range of sugars. This transporter is typically upregulated in tissue serving as a sink for photosynthate, such as roots, or as a response to mechanical damage or insect attack (Reinders et al., 2005). The trap may be absorbing prey-derived sugars or acting as a sink in the traditional sense; redistributing sugars to the trap in order to power the process of prey digestion. AKINBETA1 (AT5G21170) is also upregulated here, as it was at the 60m timepoint, implicating its importance in nutrient acquisition throughout the cycle.

Unlike the one-hour time point, at 24 hours, transcripts coding for enzymes directly related to digestion have significantly higher abundance in traps with prey: three serine-type carboxypeptidases (AT3G10410, AT1G15000, AT2G27920) two aspartic-type endopeptidases (AT3G25700 and AT2G03200), and a cysteine endopeptidase (AT5G45890), all known to be associated with the *Dionaea* secretome (Schulze et al., 2012).

The two genes with higher expression in no-prey traps at the 24-hour time point include a gene homologous to CONSTANS, the circadian clock-regulated transcription factor controlling flowering in response to changes in day length, and a basic leucine zipper transcription factor involved in anthocyanin accumulation, hypocotyl elongation, and ABA response. While CONSTANS is best known for its role in regulating flowering (Putterill et al., 1995), a CONSTANS-like gene in rice, Ghd2, was found to activate the expression of genes related to leaf senescence (Liu et al., 2016). Future functional investigations of these genes and their relationship to botanical carnivory are warranted. Taken together, we can infer based on the transcriptomic data presented here that transcriptional initiation of the digestion process does not occur before five minutes post prey capture. By 60 minutes, however, traps with prey are primed for digestion and nutrient absorption, and traps without prey have begun to repair damaged tissue ahead of trap resetting. By 24 hours, the traps with prey also begin upregulating enzyme-coding transcripts related to digestion in addition to the continuation of defense response, response to pathogens, and the production of nutrient transporters. At the same time, traps triggered without prey show the differential expression of very few genes although the trap remains closed – suggesting a refractory period before the trap resetting.

### Time series analysis

Generally speaking, our time structured experiment reveals that the process of prey capture, digestion, and nutrient absorption is a dynamic and complex process involving the transcriptional orchestration of roughly 50% of protein-coding genes in *Dionaea muscipula*. Module 3 is the most gene rich module (n = 3009) and is the only module whose peak expression is seen at the end of our time series (72 hours). This module also contains an abundance of enzymes related to prey digestion (Fig.3b)), suggesting that the digestive process remains dynamic even after several days of activity. In fact, this late peaking module is enriched in chitinase activity, amylase activity, and aspartic endopeptidase activity when compared to the modules with peak expression within a few hours of prey capture (Figure 3c). The enrichment of transcripts for some of these enzyme classes directly related to digestion suggest that this peak of expression isn’t simply related to the consequences of digestion (e.g. cell repair or senescence), but may be indicating a changing cocktail of trap fluid proteins with prey digestion and nutrient absorption functions. Another possibility is that the concentrations of digestive enzymes peak earlier than 72 hours, but the transcripts included in module 3 continue to accumulate. Further work on the dynamics of prey digestion and nutrient absorption in traps should integrate proteomic and metabolomic data.

### Conclusion

Comparisons of transcript profiles for triggered *Dionaea muscipula* traps with and without prey suggest that plants decide somewhere between the five- and sixty-minute timepoints whether they have captured suitable prey and alter their transcriptome accordingly. Differences in the transcription profiles of traps with and without prey emerge well before the trap hermetically seals and, of course, before the secretion of digestive enzymes. The upregulation of several key genes related to nutrient absorption and transport occurred early and preceded the upregulation of some of the characteristic digestive enzymes, such as the various endopeptidases. This observation suggests that prey-derived macromolecules do not need to be liberated to initiate the absorptive function of traps. Many transcripts show peak abundance within an hour of prey capture, but more than 3000 genes exhibit increasing transcript abundance up to and likely beyond 72 hours post-capture including an enrichment of amylase activity, aspartic endopeptidase activity, and chitinase activity. This observation suggests a dynamic and robust transcriptomic response related to prey digestion and nutrient absorption that continues up to and likely after three days after prey capture. In addition to integrating transcriptomic, proteomic and metabolomic analyses, better understanding of Venus flytrap prey digestions and nutrient absorption dynamics could come with increased timepoints in a time series analysis, extending beyond 72 hours.

## 6 Author Contributions

JR and JL-M conceived the study. SB performed analyses. JR and SB composed the initial draft of the manuscript and JR, SB and JL-M refined the manuscript before submission. All authors read and approved the final manuscript.

## 8 Acknowledgments

We thank Ingrid Jordan Thaden for her thoughtful advice on RNA isolation techniques and Michael Kane for *Dionaea muscipula* tissue culture. We thank Jane Grimwood, Director of the Genome Sequencing Center (GSC) at HudsonAlpha Institute for Biotechnology, for providing outstanding sequencing services. We also thank the systems administrators and support staff at the Georgia Advanced Computing Resource Center (GACRC) for maintaining the high-performance computing platforms used for all our bioinformatic analyses.

## 9 Data Availability Statement

Scripts and results for these analyses can be found at: https://github.com/mellamosummer/2022VenusFlyTrap/tree/main. Reads have been deposited in NCBI’s SRA database under the project ID PRJNA981540

## 10 Supplementary Material

A) BLAST results for Dm Transcripts B) 1 hr prey/noprey DEG list C) 24 hr prey/no prey DEG list D) GO MF Enrichment for 1 hr prey/no prey DEG list E) GO MF Enrichment for 24 hr prey/noprey DEG list F) GO BP Enrichment for 1 hr prey/no prey DEG list G) GO BP Enrichment for 24 hr prey/no prey DEG list H) Transcripts per million across libraries for Gene Coexpression Analysis I) Library Metadata for Gene Coexpression Analysis J) Network Modules K) GO MF Enrichment for Module 2 L) GO MF Enrichment for Module 3 M) GO MF Enrichment for Module 4 N) GO MF Enrichment for Module 6 O) GO MF Enrichment for Module 7 P) GO MF Enrichment for Module 9 Q) GO MF Enrichment for Module 10 R) GO MF Enrichment for Module 16 S) GO BP Enrichment for Module 2 T) GO BP Enrichment for Module 3 U) GO BP Enrichment for Module 4 V) GO BP Enrichment for Module 6 W) GO BP Enrichment for Module 7 X) GO BP Enrichment for Module 9 Y) GO BP Enrichment for Module 10 Z) GO BP Enrichment for Module 16 AA) QC AB) Libraries and Sequencing AC) Read Mapping

